# Fractal dimension of EEG signal senses complexity of fractal animations

**DOI:** 10.1101/2021.02.11.430870

**Authors:** Sarshar Dorosti, Reza Khosrowabadi

## Abstract

We are surrounded with many fractal and self-similar patterns which has been area of many researches in the recent years. We can perceive self-similarities in various spatial and temporal scales; however, the underlying neural mechanism needs to be well understood. In this study, we hypothesized that complexity of visual stimuli directly influence complexity of information processing in the brain. Therefore, changes in fractal pattern of EEG signal must follow change in fractal dimension of animation. To investigate this hypothesis, we recorded EEG signal of fifteen healthy participants while they were exposed to several 2D fractal animations. Fractal dimension of each frame of the animation was estimated by box counting method. Subsequently, fractal dimensions of 32 EEG channels were estimated in a frequency specific manner. Then, association between pattern of fractal dimensions of the animations and pattern of fractal dimensions of EEG signals were calculated using the Pearson’s correlation algorithm. The results indicated that fractal animation complexity is mainly sensed by changes in fractal dimension of EEG signals at the centro-parietal and parietal regions. It may indicate that when the complexity of visual stimuli increases the mechanism of information processing in the brain also enhances its complexity to better attend and comprehend the stimuli.

## Introduction

Visual sense has an important role in shaping human understanding of the natural world. Therefore, the way that brain is involved in perception of natural world has always been of interest to researchers. Many studies have been conducted on understanding effects of the texture, color, light and frequency of visual stimuli. In this regard, various neuroimaging techniques have been implied to determine underlying brain mechanism while ones process effects of each of them (Yoto, Katsuura et al. 2007, Barrett, Beaulieu et al. 2010, Vurro, Ling et al. 2013). Among the neuroimaging techniques, electroencephalogram (EEG) is relatively more accessible and could map the electrical activities of the brain with an accurate time resolution. Hence, many EEG studies have been performed and several mathematical and computational methods have been used to investigate characteristics of EEG depends on the type of external stimulus (Isaksson, Wennberg et al. 1981, Srinivasan 2007, Kim, Kim et al. 2013, Namazi 2018). For instance, fractal analysis and employed approximate entropy have been implied to talk about randomness and complexity of EEG signal (Namazi, Akrami et al. 2016). Complexity of EEG signal measured by fractal dimension (FD) presents self-similarities across different scales (Mandelbrot 1982).

Fractal concept can describe inherent irregularity and complexity of natural objects, such as scaling properties of the EEG signal. So, fractal dimension is viewed as an index of complexity, which shows how a detail in a pattern changes with a scale at which it is measured. Fractal approach has been used widely to study the complexity of different biomedical signal and patterns such as DNA (Glenny, Robertson et al. 1991, Arneodo, d’Aubenton-Carafa et al. 1996, Zu-Guo, Anh et al. 2002, Butala and Sadana 2003, Cattani 2010, Namazi and Kiminezhadmalaie 2015), human face (Namazi, Akrami et al. 2016), respiration signal (Peng, Mietus et al. 2002), eye movement time series (Taylor, Micolich et al. 1999, Alipour, Namazi et al. 2019), cell image analysis (Smith Jr, Marks et al. 1989), roughness of texture (Chappard, Degasne et al. 2003), bone structure (Jennane, Ohley et al. 2001), heart rate (Mäkikallio, Huikuri et al. 2001), spider brain signal (Namazi 2017), and movement behavior of animal in foraging (Namazi 2017). Likewise, there are several reported works that employed fractal dimension for analysis of EEG signal (Lutzenberger, Preissl et al. 1995, Accardo, Affinito et al. 1997, Acharya, Faust et al. 2005, Harne 2014, Klonowski 2016). For instance, studies on employing fractal theory in analysis of EEG signal during process of visual (Namazi, Kulish et al. 2016, Namazi, Ala et al. 2018, Alipour, Towhidkhah et al. 2019), auditory (Namazi, Khosrowabadi et al. 2016, Reza Namazi 2017, Alipour, Khosrowabadi et al. 2018) and olfactory (Omam, Babini et al. 2020) stimuli, in healthy and patient subjects are noteworthy to be mentioned. It should be noted that in all the reported studies, scientists found variations of fractal structure in EEG signal due to external stimulation. Nevertheless, it is still not clear that how complexity of information in visual stimuli influence the complexity of EEG signal.

Considering the complex fractal phenomena in our peripheral environment, in this study, we hypothesized that complexity of visual stimuli linearly influence the complexity of EEG signals as outcome of information processing in the brain. Our aim is to show fractal patterns of brain electrical activities are positively correlated with fractal patterns of exposed visual stimuli. Therefore, we selected a group of healthy volunteers and exposed them to some 2 and 3 dimensional (2D and 3D) fractal animations while their brain activities were recorded. Subsequently, correlation between FDs of the animations and FDs of EEG signals was calculated. Methodology, experimental results and discussion about the findings are presented in the following sections.

## Materials and methods

### Participants

In this paper, it is intended to perform fractal analysis on the fractal animations and their correspondent EEG data recorded from young and healthy individuals. Therefore, a group of 15 healthy postgraduate university students (all male, 25.93 ±1.79 years old, 3 left-handed) were recruited. The sample size was suggested by the GPower framework considering p value of 0.05 for pearson’s correlation was 12, that considering 25% drop-out, 15 subjects were recruited. Subjects were non-smokers and did not have any family history of mental disorder. They were asked to not take any medicine during last 24 hours, and not drink coffee 4 hours before the experiment. All the participants filled a consent form prior to the experiment. The study was part of a bigger project approved by the Iran Medical School review board in accordance with the Helsinki declaration (IR.IUMS.REC.1395.933091506).

### Experimental design

Experimental paradigm and visual stimuli implied in this study are illustrated in Figue 1 and Table 1.

**Table 1.**
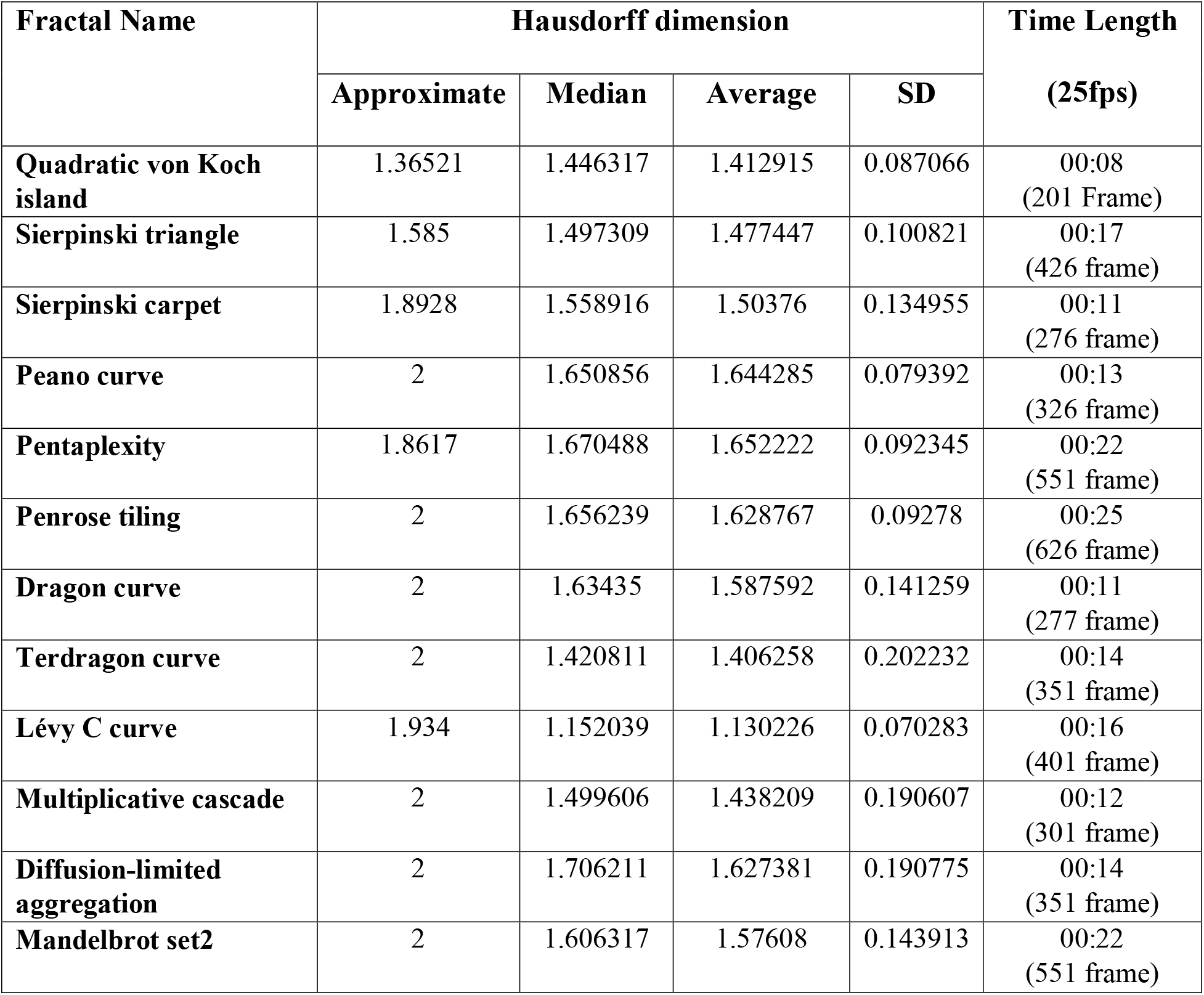
Characteristics of implied fractal animations

**Figure 1.**
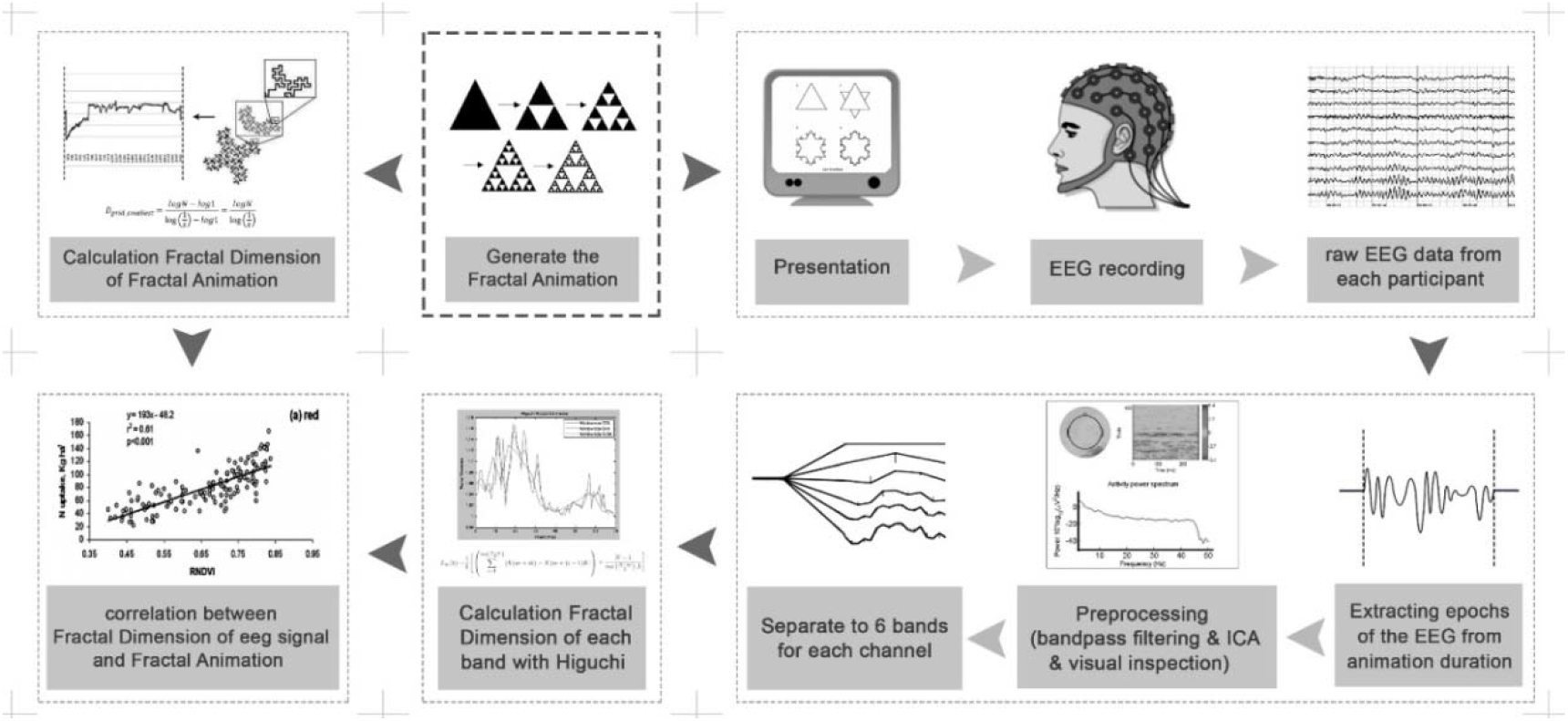
Schematic of experimental paradigm and data analysis procedure

#### Visual stimuli: Fractal animations

As indicated in Table1, 12 types of fractal animations were generated as visual stimuli in 2D and 3D shapes. The current study is only based on the Timing of each animation was in range of 10 to 60 seconds (24 frames per second). Additionally, after visual inspection we applied edge detection algorithm to all the frames in video processing level. Then, we used Box-counting method (Sarkar and Chaudhuri 1994, Raghavendra and Dutt 2010) in TOUCHDESIGNER 0.99 (derivative.ca) and PYTHON 3.5 (www.python.org) to calculate the FDs of all animation frames. The 3D fractal animations were also generated with Mandelbulb 3D (MB3D) 1.9.1 software (www.mandelbulb.com).

Subsequently, visual stimuli were presented on a 42" monitor placed 1.3 m away from the subjects. The visual stimuli were presented to the subjects while their EEG data was collected. The raw data for this work was collected at the Institute for Cognitive and Brain Sciences (ICBS), Tehran in late spring and early summer of 2019. To reduce physiological variability, the EEG data were only recorded from 9 to 11 AM at a dim-light, silent and temperature controlled room. The EEG data were recorded by an EEG8 amplifier with sampling frequency of 1024 (Contact Precision Instruments, Cambridge, MA) use of 32 electrodes (Electrocap International, Inc) placed in the 10-20 international standard positions. Synchronization between visual stimuli presentation and EEG data recording was performed by sending trigger markers through the parallel port using the Psychtoolbox 3.0.13 in MATLAB (The MathWorks Inc, Natick, USA).

### Data analysis

#### Fractal dimensions of visual stimuli

FD is an important feature of fractal geometry which has many applications in various fields including image processing, image analysis, texture segmentation, shape classification and identifying the image features including roughness and smoothness. The fundamental rule of computing fractal dimension of a whole image depends upon the theory of self-similarity. Self-similarity is significant assets of fractal theory which basically was used to calculate the FD. There are many techniques to estimate dimension of a fractal surface. One of the most famous technique is the grid dimension method which is popularly known as box counting (Sarkar and Chaudhuri 1994, Raghavendra and Dutt 2010). The box-counting method and its derivatives including differential box counting and improved differential box counting (Sarkar and Chaudhuri 1994) methods have been used to estimate FD of grayscale images. It should be mentioned that color images are generally converted to grayscale and their FDs or their roughness are then calculated. (Panigrahy, Seal et al. 2019)

#### Fractal dimension of EEG data

Regarding the neurophysiological data, a standard preprocessing pipeline was performed on the raw EEG data. The EEG data was down-sampled to 512 Hz and using a simple FIR filter with zero phase-shift, the EEG data were band-pass filtered between 1 to 40Hz. After segmenting the whole data to 3-second trials, artifacts were removed from the data using the ICA technique, followed by a visual inspection. Bad channels were detected using the kurtosis method, and were interpolated to the average of their neighbors’ activities. Finally, EEG data were re-referenced to average activities of all the channels. The preprocessing was performed using MATLAB R2016b (The MathWorks Inc., Natick, USA) and the EEGLAB v14.1.2b toolbox.

Subsequently, since the brain works in a self-organized manner described by specific frequency bands, the EEG data were filtered to 5 conventional frequency bands including delta [1-4Hz], theta [4-8 Hz], Alpha [8-13 Hz], Beta [13-30 Hz] and lower Gamma [30-40 Hz]. Then, the fractal dimensions of EEG data were calculated at each frequency band separately. The FD of preprocessed EEG signals from 15 healthy controls were separately computed using Higuchi’s method (Higuchi 1988).

After calculation of FDs of animations and fractal dimensions of EEG data, association between FDs of EEG data and fractal dimensions of animations were estimated using the Pearson correlation coefficient. Threshold of 0.05 was considered for the significance of the results.

## Results

A fractal is a mathematical set that showcases a recurrent pattern in every increasingly small scale (Mandelbrot and Aizenman 1979). Fractal dimension calculates the effective number of degrees of freedom in a dynamical system (Mandelbrot 1982) and is a ratio that represents a statistical index of complexity(Theiler 1990). The box counting method is widely used for estimation of fractal dimension since it is ease to implement and provide an accurate estimation [(Kyriacos, Buczkowski et al. 1994, Li, Du et al. 2009)]. The box counting algorithm finds an optimized way of cutting a pattern up and reveals the scaling factor. Therefore, we processed complexity of each frame of animations using the box-counting algorithm.

Figure 2 presents an example pattern of changes in the fractal dimension of a visual stimulus (fractal animation).

**Figure 2.**
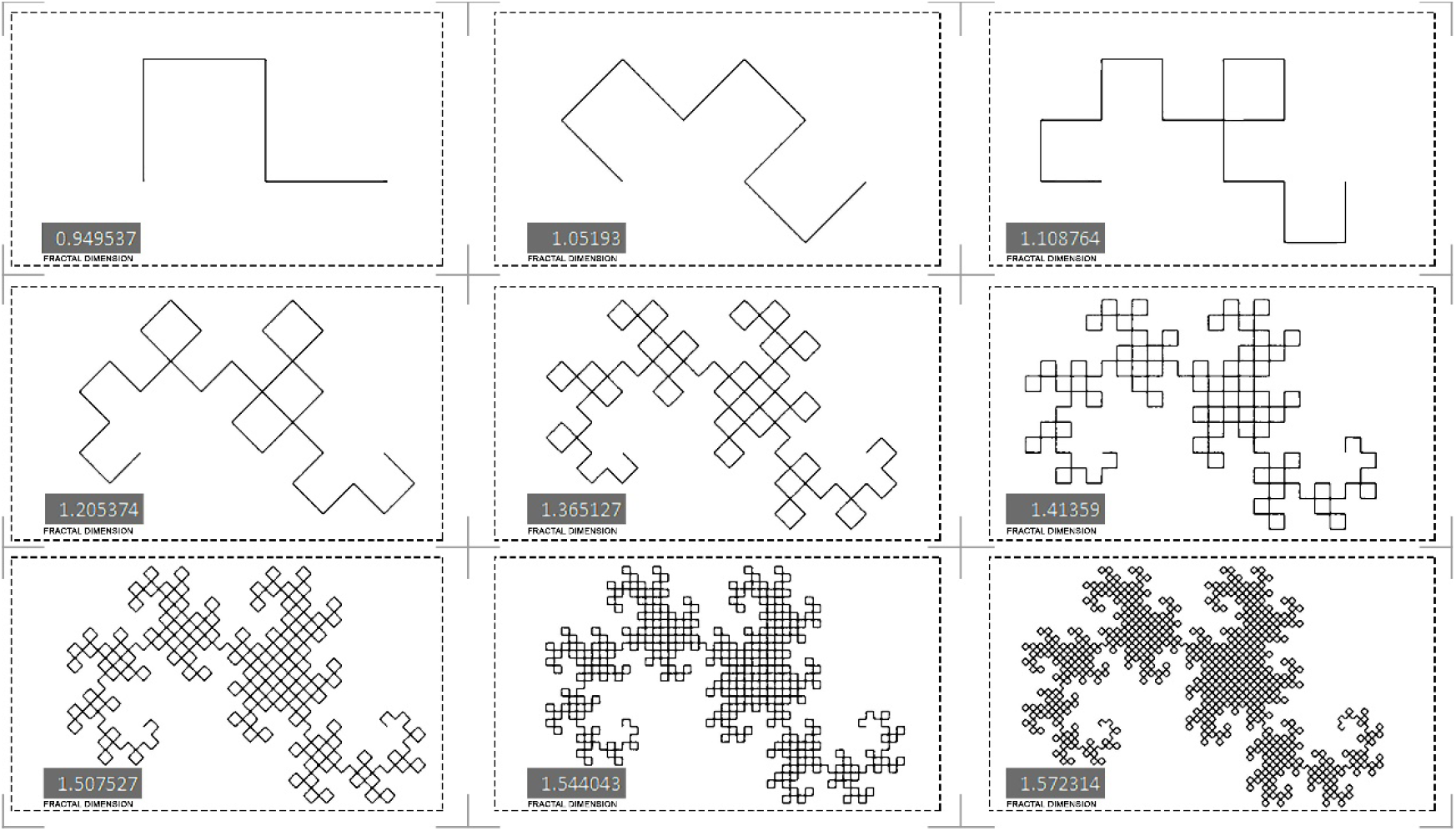
FD changes in Terdragon curve animation. The number at the bottom left corner denotes the fractal dimension of the image (one frame of animation).

We designed changes in the complexity of the fractal animations to be unpredictable and complete data of fractal animations are provided in the supplementary data. Time series of FD changes in one of the animations is presented in Figure 3.

**Figure 3.**
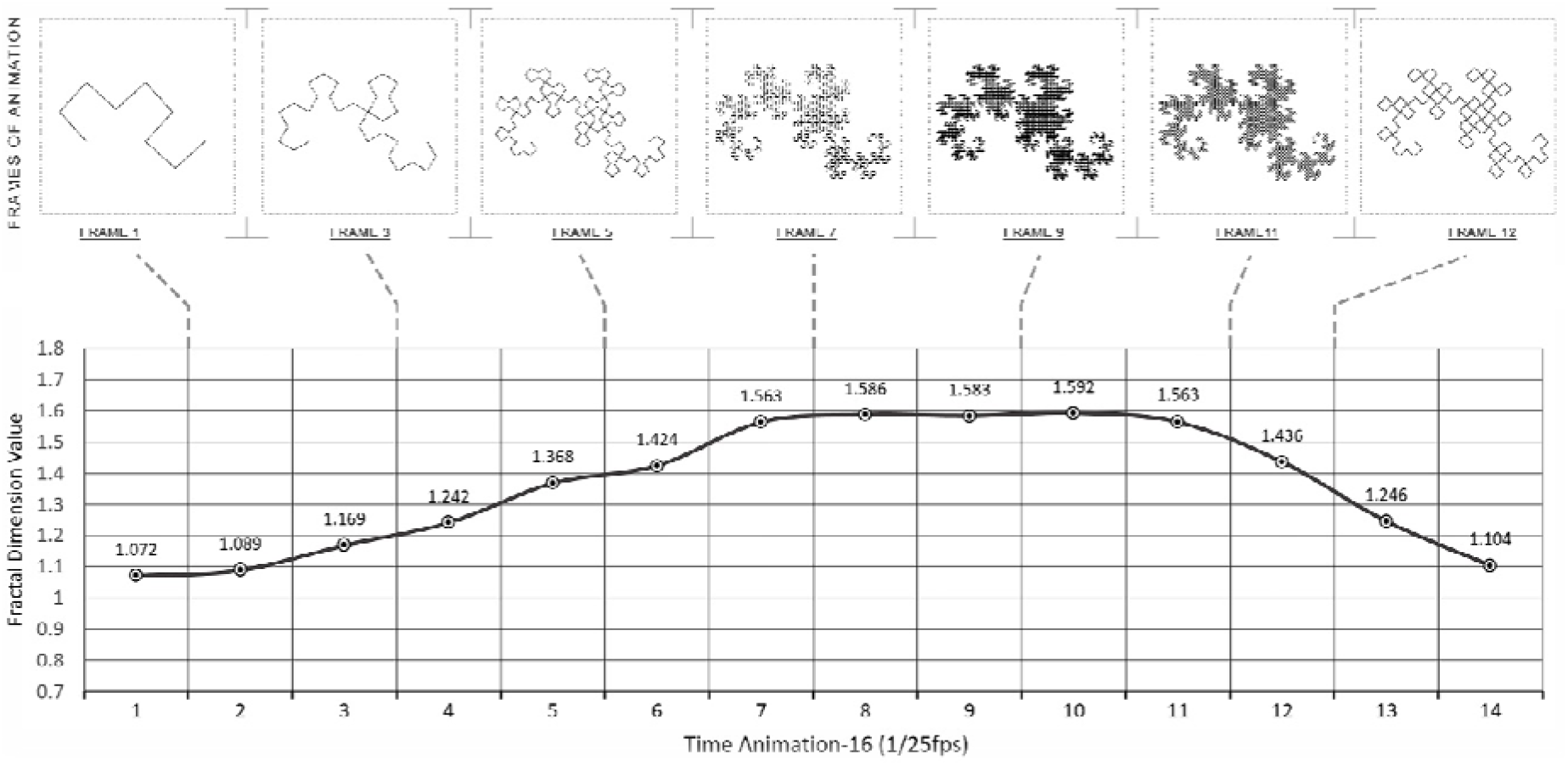
Pattern of changes in fractal dimension of Terdragon curve animation. A: FD of each frame image of animation. B: Series of changes in FDs of animation frames.

In addition, there are several approaches to compute fractal dimension of EEG time series, including: Higuchi’s method (Higuchi 1988), Katz’s method(Katz 1988), box-counting (Sarkar and Chaudhuri 1994, Raghavendra and Dutt 2010). There are studies that have compared performances of the aforementioned algorithms in terms of accuracy, sensitivity to the sampling frequency and their dependency of the estimation to the selected length of time window (Raghavendra and Dutt 2010). It is proposed that the Higuchi’s method of computation of FD is the robust and gives accurate estimation results (Accardo, Affinito et al. 1997, Raghavendra and Dutt 2010) Therefore we implied the higuchi method for FD of EEG that time series of cganges in FDs of EEG are presented at the Figure 4.

**Figure 4.**
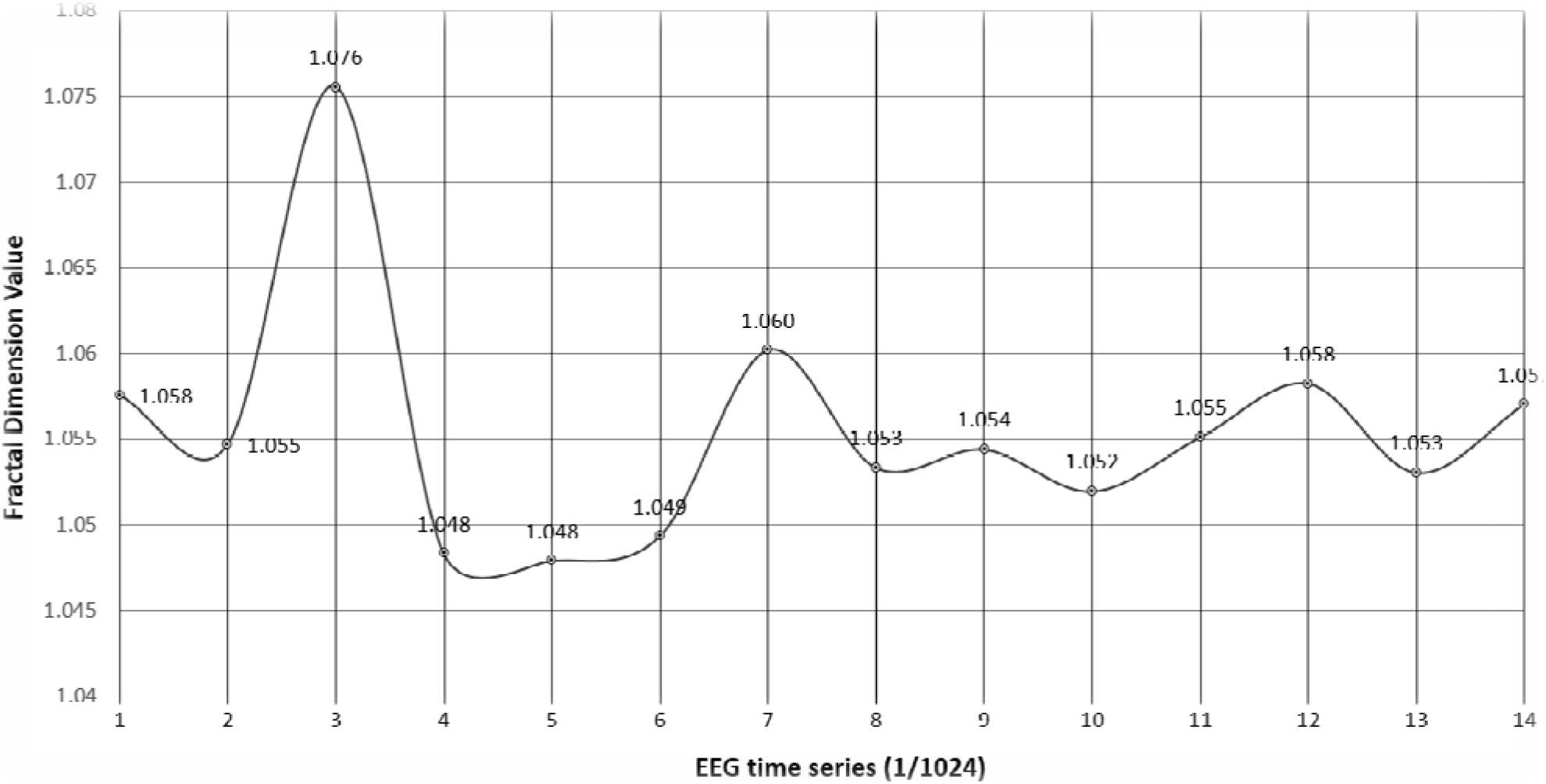
Series of changes in fractal dimension of EEG signal while watching Terdragon curve animation

Significant correlation between FDs of fractal animations and FDs of EEG data were identified at theta at the frontal regions, and at the alpha, and the beta bands in central and parietal regions. Figure 5 illustrates the topoplots of average of Pearson’s correlation values observed in all the subjects in a band-specific manner. The similar procedure was performed for all the animations which the results are separately presented in supplementary material.

**Figure 5.**
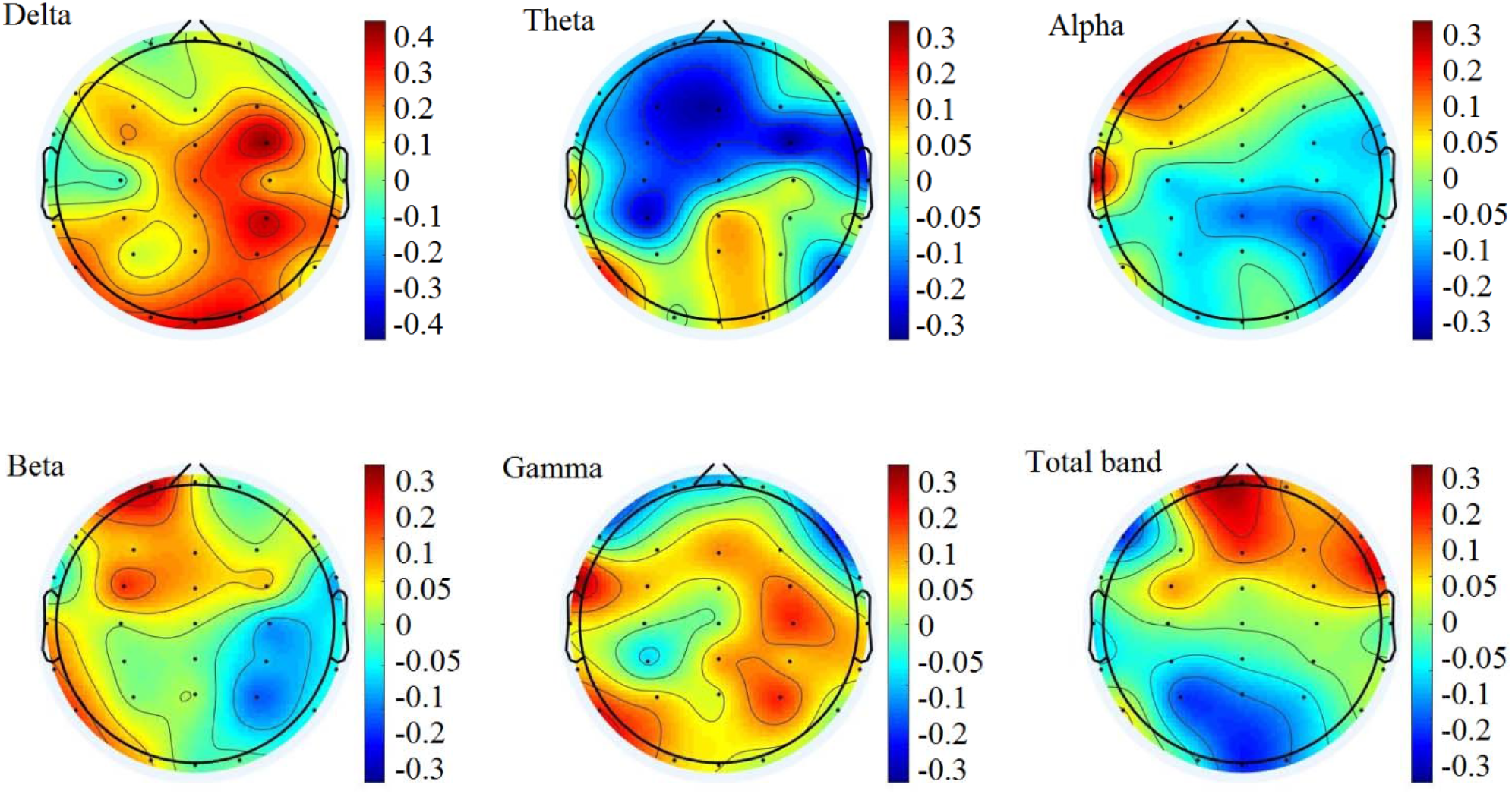
Pearson’s correlation between series of FDs of the Terdragon curve animation and series of the correspondent EEG FDs. The color bars indicate average of correlation values in all the subjects.

FDs of EEG signals mainly at the centro-parietal and parietal channels including Pz, Cpz, Cp3 showed correlation with FDs of 2D animations. In addition, frequency-band specific analysis also showed significant correlation at the P3, Pz, Cp3, Cpz, C3 in alpha band, and Cz, Cpz, Pz, Cp4 in beta band. The results indicated that EEG signals at the parietal regions have more complex pattern while 2D animations with higher FDs were presented.

## Discussion and conclusion

Beside all works performed in this area of research, brain mechanism underlying perception of fractal animations requires to be well understood. In this study, we hypothesized that complexity of visual stimuli directly influence complexity of information processing in the brain. Under the assumption that EEG signals show the underlying neural mechanism of perception of visual stimuli, analysis of the EEG data could help to study the dynamical properties of information processing in the brain. We used fractal dimension as a measure of complexity and calculated complexity of patterns in the frames of fractal animations with box-counting method. In addition, complexity of EEG data in healthy control subjects was also estimated using the Higuchi’s fractal dimension. Subsequently, association between time series of FDs of fractal animations and time series of EEG FDs were calculated. Our results showed complexity of visual stimuli could linearly influence complexity of information processing in the brain. The pattern of changes showed that fractal patterns of brain electrical activities mainly at the posterior part of brain were positively correlated with fractal patterns of exposed visual stimuli. It denotes that when the self-similarities in the fractal animations were increased, in the same way the self-similarities of brain signals also were increased. It means that the mechanism of information processing in the brain also enhance its complexity to better attend and comprehend the stimuli.

The method discussed in this research can be further implied on scenes with fractal properties in cinema, animation, video games or advertising industry. In addition, a mathematical model which makes a link between the fractal properties of fractal animations and EEG signals can also be used in therapeutic or commercial applications. Such a finding could help us to pave the way to better understanding the brain behavior and has a great potential to be used in brain-computer interface applications as well. Investigation on diversity of animations and other measures of complexity as well as looking at possible applications of the findings are proposed for future works.

It should be mentioned that because of some practical issues, we only used male subjects in our study. Therefore, it is impossible to apply our findings to females or to anyone outside the implied age range. In addition, the number of participants was small. So, the extension of the study framework with larger sample sizes could provide more robust results.

## Acknowledgments

We would like to thank Mr. Mohsen Shabani for his generous help in data gathering stage and all the participants who helped us in this study.

## Conflict of interest

Authors declare no conflict of interest.

## Notes

### Competing Interest Statement

The authors have declared no competing interest.

## References

Accardo, A., et al. (1997). “Use of the fractal dimension for the analysis of electroencephalographic time series.” Biological Cybernetics 77(5): 339–350.

Acharya, R., et al. (2005). “Non-linear analysis of EEG signals at various sleep stages.” Computer methods and programs in biomedicine 80(1): 37–45.

Alipour, H., et al. (2019). “Fractal-based analysis of the influence of color tonality on human eye movements.” Fractals 27(03): 1950040.

Alipour, H., et al. (2019). “Complexity-based analysis of the relation between fractal visual stimuli and fractal eye movements.” Fluctuation and Noise Letters 18(03): 1950012.

Alipour, Z. M., et al. (2018). “Fractal-based analysis of the influence of variations of rhythmic patterns of music on human brain response.” Fractals 26(05): 1850080.

Arneodo, A., et al. (1996). “Wavelet based fractal analysis of DNA sequences.” Physica D: Nonlinear Phenomena 96(1-4): 291–320.

Barrett, D. M., et al. (2010). “Color, flavor, texture, and nutritional quality of fresh-cut fruits and vegetables: desirable levels, instrumental and sensory measurement, and the effects of processing.” Critical reviews in food science and nutrition 50(5): 369–389.

Butala, H. D. and A. Sadana (2003). “A fractal analysis of analyte–estrogen receptor binding and dissociation kinetics using biosensors: environmental effects.” Journal of colloid and interface science 263(2): 420–431.

Cattani, C. (2010). “Fractals and hidden symmetries in DNA.” Mathematical problems in engineering 2010.

Chappard, D., et al. (2003). “Image analysis measurements of roughness by texture and fractal analysis correlate with contact profilometry.” Biomaterials 24(8): 1399–1407.

Glenny, R. W., et al. (1991). “Applications of fractal analysis to physiology.” Journal of Applied Physiology 70(6): 2351–2367.

Harne, B. P. (2014). “Higuchi Fractal Dimension Analysis of EEG Signal before and after OM Chanting to Observe Overall Effect on Brain.” International Journal of Electrical & Computer Engineering (2088-8708) 4(4).

Higuchi, T. (1988). “Approach to an irregular time series on the basis of the fractal theory.” Physica D: Nonlinear Phenomena 31(2): 277–283.

Isaksson, A., et al. (1981). “Computer analysis of EEG signals with parametric models.” Proceedings of the IEEE 69(4): 451–461.

Jennane, R., et al. (2001). “Fractal analysis of bone X-ray tomographic microscopy projections.” IEEE transactions on medical imaging 20(5): 443–449.

Katz, M. J. (1988). “Fractals and the analysis of waveforms.” Computers in biology and medicine 18(3): 145–156.

Kim, M.-K., et al. (2013). “A review on the computational methods for emotional state estimation from the human EEG.” Computational and mathematical methods in medicine 2013.

Klonowski, W. (2016). Fractal Analysis of Electroencephalographic Time Series (EEG Signals). The Fractal Geometry of the Brain, Springer: 413–429.

Kyriacos, S., et al. (1994). “A modified box-counting method.” Fractals 2(02): 321–324.

Li, J., et al. (2009). “An improved box-counting method for image fractal dimension estimation.” Pattern Recognition 42(11): 2460–2469.

Lutzenberger, W., et al. (1995). “Fractal dimension of electroencephalographic time series and underlying brain processes.” Biological Cybernetics 73(5): 477–482.

Mäkikallio, T. H., et al. (2001). “Fractal analysis and time-and frequency-domain measures of heart rate variability as predictors of mortality in patients with heart failure.” The American journal of cardiology 87(2): 178–182.

Mandelbrot, B. and M. Aizenman (1979). “Fractals: form, chance, and dimension.” PhT 32(5): 65.

Mandelbrot, B. B. (1982). “The Fractal Geometry of.” Nature: 394–397.

Namazi, H. (2018). “Complexity based analysis of the correlation between external stimuli and bio signals.” ARC J. Neurosci. 3(3): 6–9.

Namazi, H., et al. (2016). “The fractal based analysis of human face and DNA variations during aging.” BioScience Trends.

Namazi, H., et al. (2016). “Analysis of the influence of complexity and entropy of odorant on fractal dynamics and entropy of EEG signal.” BioMed Research International 2016.

Namazi, H., et al. (2018). “Decoding of steady-state visual evoked potentials by fractal analysis of the electroencephalographic (EEG) signal.” Fractals 26(06): 1850092.

Namazi, H., et al. (2016). “Analysis of the influence of memory content of auditory stimuli on the memory content of EEG signal.” Oncotarget 7(35): 56120.

Namazi, H. and M. Kiminezhadmalaie (2015). “Diagnosis of lung cancer by fractal analysis of damaged DNA.” Computational and mathematical methods in medicine 2015.

Namazi, H., et al. (2016). “The analysis of the influence of fractal structure of stimuli on fractal dynamics in fixational eye movements and EEG signal.” Scientific reports 6: 26639.

Namazi, H. R. (2017). “The complexity based analysis of the correlation between spider’s brain signal and web.” ARC Journal of Neuroscience 2(4): 38–44.

Namazi, H. R. (2017). “Fractal based analysis of movement behavior in animal foraging.” ARC J. Neurosci. 2(3): 1–3.

Omam, S., et al. (2020). “Complexity-based decoding of brain-skin relation in response to olfactory stimuli.” Computer methods and programs in biomedicine 184: 105293.

Panigrahy, C., et al. (2019). “Differential box counting methods for estimating fractal dimension of gray-scale images: A survey.” Chaos, Solitons & Fractals 126: 178–202.

Peng, C.-K., et al. (2002). “Quantifying fractal dynamics of human respiration: age and gender effects.” Annals of biomedical engineering 30(5): 683–692.

Raghavendra, B. and D. N. Dutt (2010). “Computing fractal dimension of signals using multiresolution box-counting method.” International Journal of Information and Mathematical Sciences 6(1): 50–65.

Reza Namazi, H. (2017). “Fractal-based analysis of the influence of music on human respiration.” Fractals 25(06): 1750059.

Sarkar, N. and B. B. Chaudhuri (1994). “An efficient differential box-counting approach to compute fractal dimension of image.” IEEE Transactions on systems, man, and cybernetics 24(1): 115–120.

Smith Jr, T., et al. (1989). “A fractal analysis of cell images.” Journal of neuroscience methods 27(2): 173–180.

Srinivasan, N. (2007). “Cognitive neuroscience of creativity: EEG based approaches.” Methods 42(1): 109–116.

Taylor, R. P., et al. (1999). “Fractal analysis of Pollock’s drip paintings.” Nature 399(6735): 422–422.

Theiler, J. (1990). “Estimating fractal dimension.” JOSA A 7(6): 1055–1073.

Vurro, M., et al. (2013). “Memory color of natural familiar objects: Effects of surface texture and 3-D shape.” Journal of Vision 13(7): 20–20.

Yoto, A., et al. (2007). “Effects of object color stimuli on human brain activities in perception and attention referred to EEG alpha band response.” Journal of physiological anthropology 26(3): 373–379.

Zu-Guo, Y., et al. (2002). “Fractals in DNA sequence analysis.” Chinese Physics 11(12): 1313.

