## Supplementary material for "Fractal dimension of EEG signal senses complexity of fractal animations": Table1

**Table1.** Characteristics of implied fractal animations

| **Fractal Name** | **Hausdorff dimension** | | | | **Time Length**  **(25fps)** |
| --- | --- | --- | --- | --- | --- |
|  | **Approximate** | **Median** | **Average** | **SD** |  |
| **Quadratic von Koch island** | 1.36521 | 1.446317 | 1.412915 | 0.087066 | 00:08  (201 Frame) |
| **Sierpinski triangle** | 1.585 | 1.497309 | 1.477447 | 0.100821 | 00:17  (426 frame) |
| **Sierpinski carpet** | 1.8928 | 1.558916 | 1.50376 | 0.134955 | 00:11  (276 frame) |
| **Peano curve** | 2 | 1.650856 | 1.644285 | 0.079392 | 00:13  (326 frame) |
| **Pentaplexity** | 1.8617 | 1.670488 | 1.652222 | 0.092345 | 00:22  (551 frame) |
| **Penrose tiling** | 2 | 1.656239 | 1.628767 | 0.09278 | 00:25  (626 frame) |
| **Dragon curve** | 2 | 1.63435 | 1.587592 | 0.141259 | 00:11  (277 frame) |
| **Terdragon curve** | 2 | 1.420811 | 1.406258 | 0.202232 | 00:14  (351 frame) |
| **Lévy C curve** | 1.934 | 1.152039 | 1.130226 | 0.070283 | 00:16  (401 frame) |
| **Multiplicative cascade** | 2 | 1.499606 | 1.438209 | 0.190607 | 00:12  (301 frame) |
| **Diffusion-limited aggregation** | 2 | 1.706211 | 1.627381 | 0.190775 | 00:14  (351 frame) |
| **Mandelbrot set2** | 2 | 1.606317 | 1.57608 | 0.143913 | 00:22  (551 frame) |
