## Supplementary Material for "Fractal dimension of EEG signal senses complexity of fractal animations"

**Title:**

**Running title:** Neural correlates of watching fractal animations

***Corresponding author:** Reza Khosrowabadi

**Address:** Institute for Cognitive and Brain Science, Shahid Beheshti University, Evin Sq., Tehran19839-63113, Iran

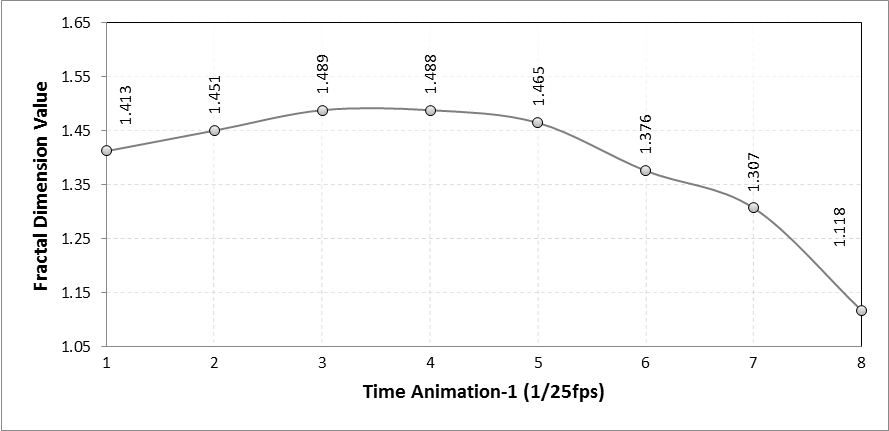

Figure1. Series of fractal dimensions of animation frames in the Quadratic von Koch island

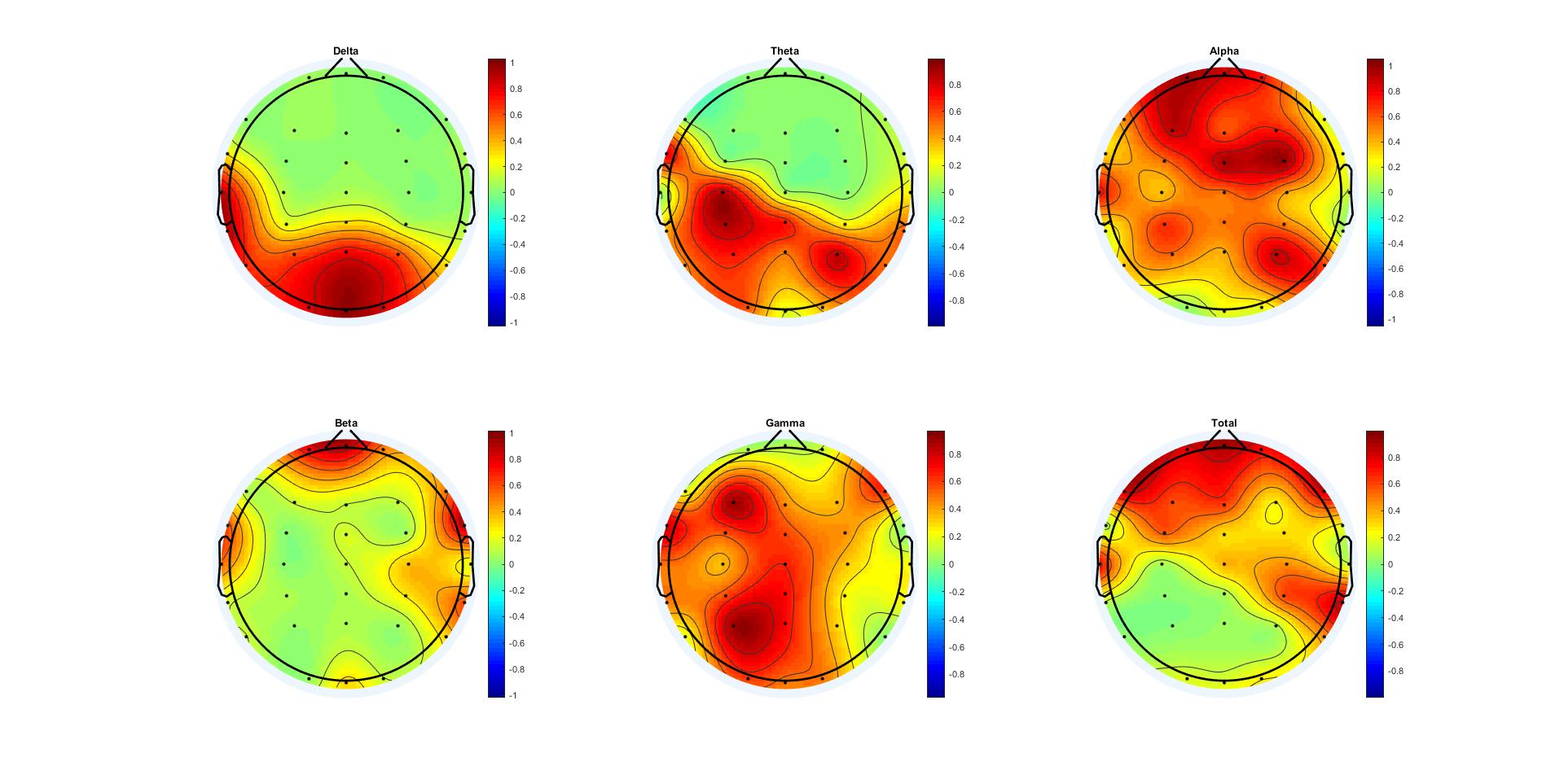

Figure2. Significant correlation between FDs of EEG signals and FDs of Quadratic von Koch island animation. Colorbars indicate the Pearson's' p value.

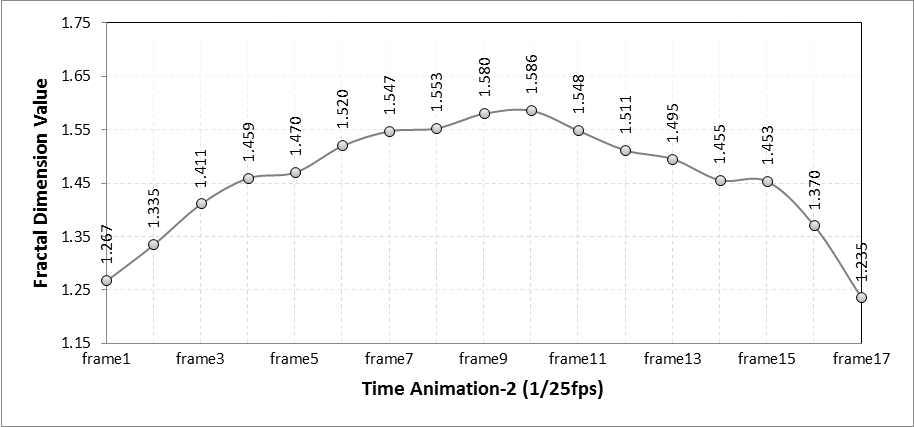

Figure3. Series of fractal dimensions of animation frames in the Sierpinski triangle

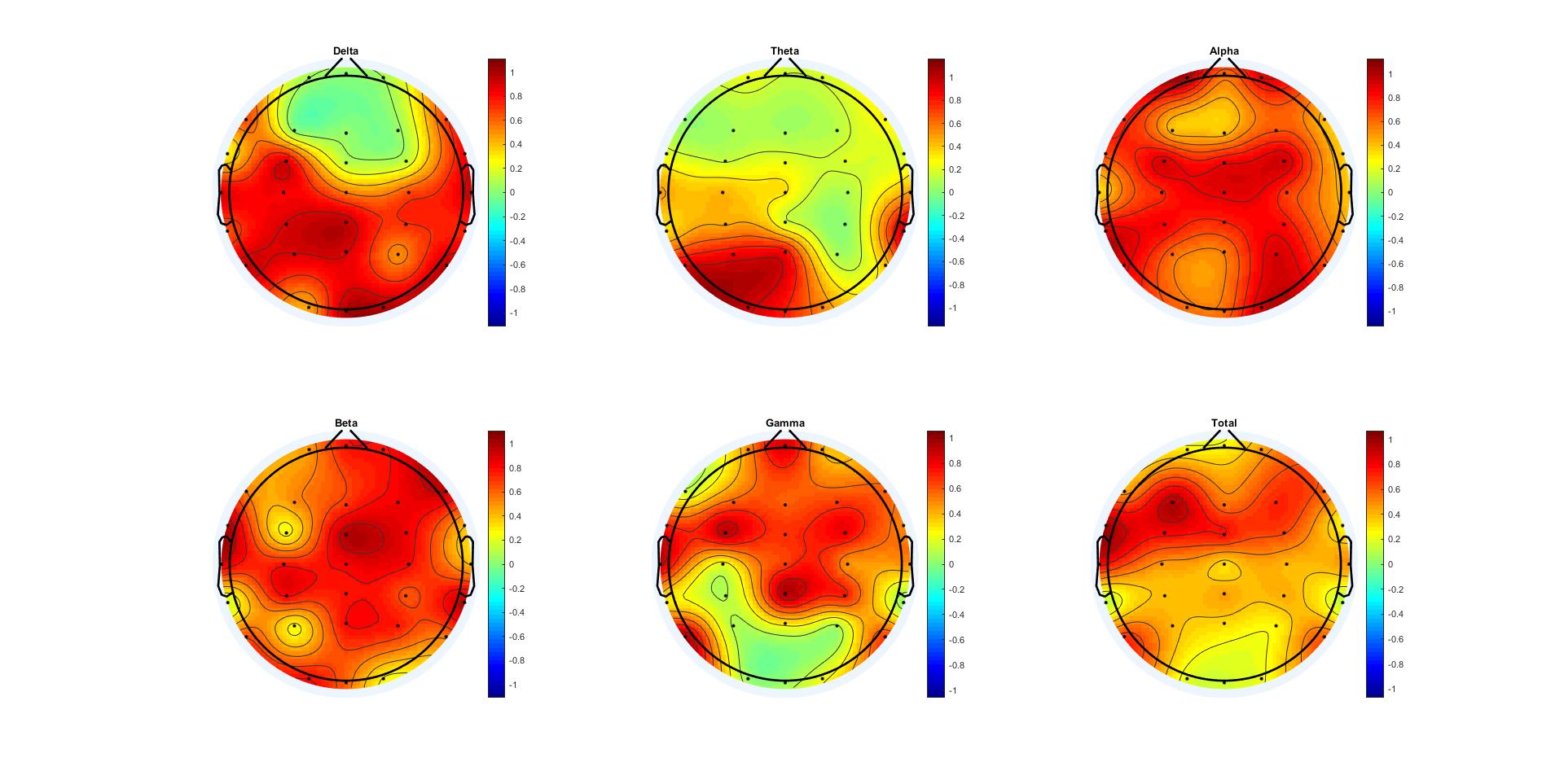

Figure4. Significant correlation between FDs of EEG signals and FDs of Sierpinski triangle animation. Colorbars indicate the Pearson's' p value.

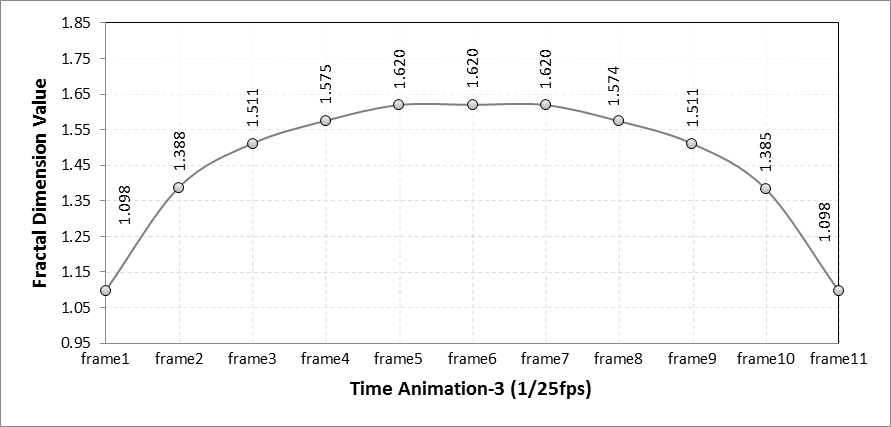

Figure5. Series of fractal dimensions of animation frames in the Sierpinski carpet

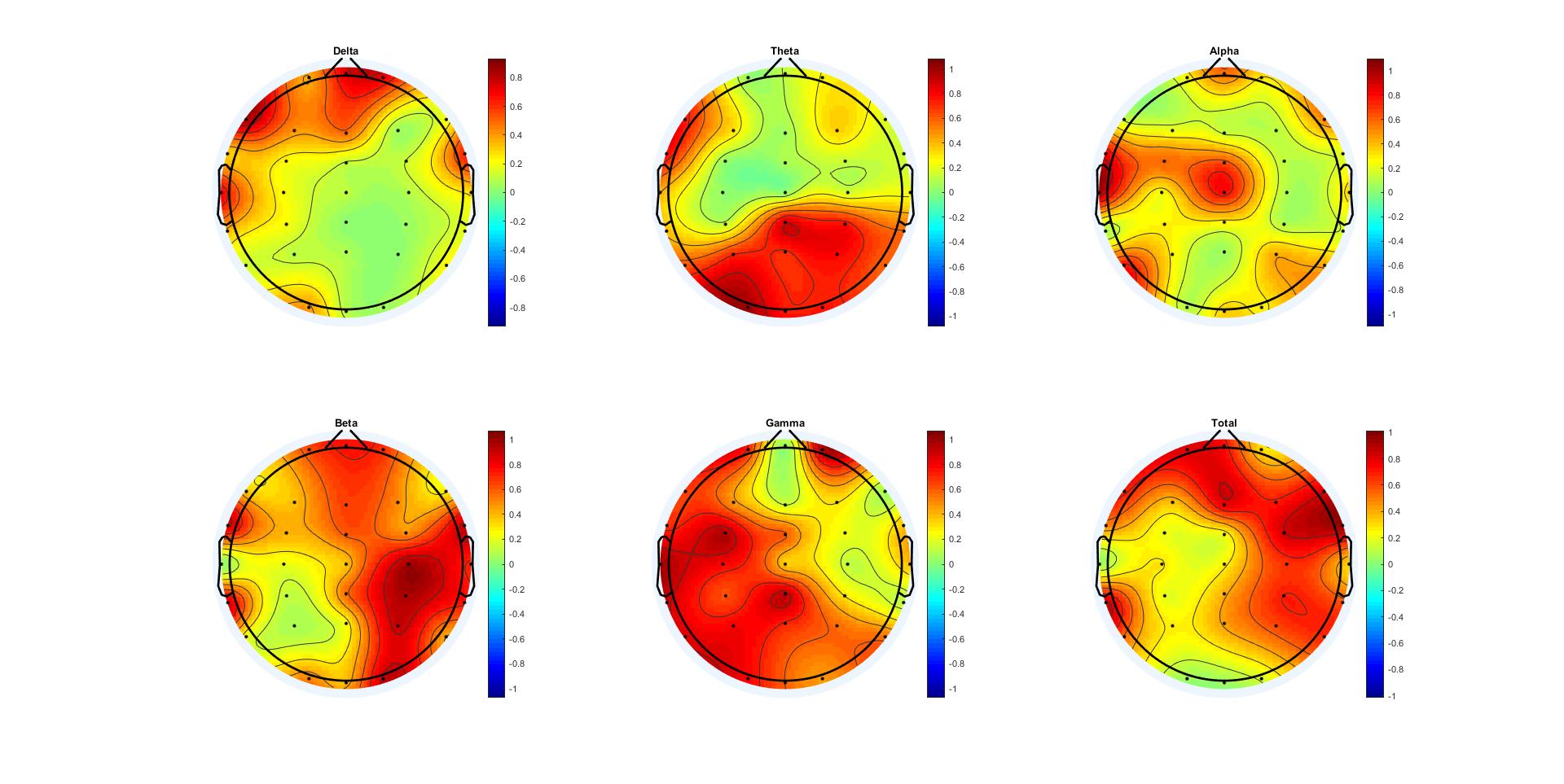

Figure6. Significant correlation between FDs of EEG signals and FDs of Sierpinski carpet animation. Colorbars indicate the Pearson's' p value.

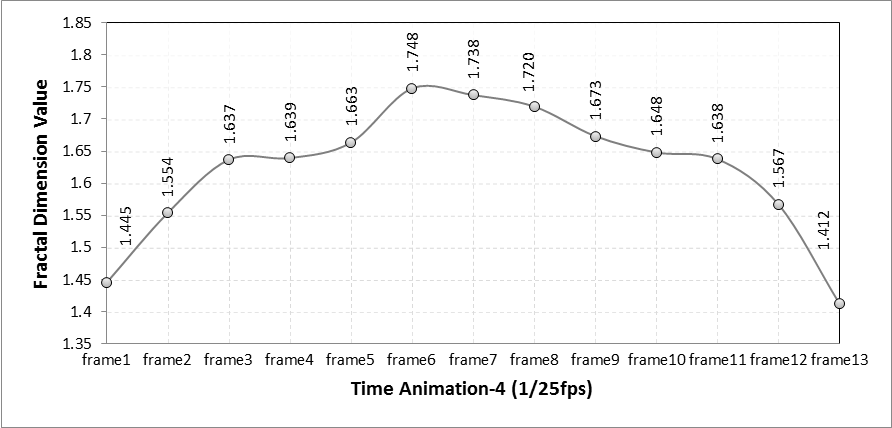

Figure7. Series of fractal dimensions of animation frames in the Peano curve

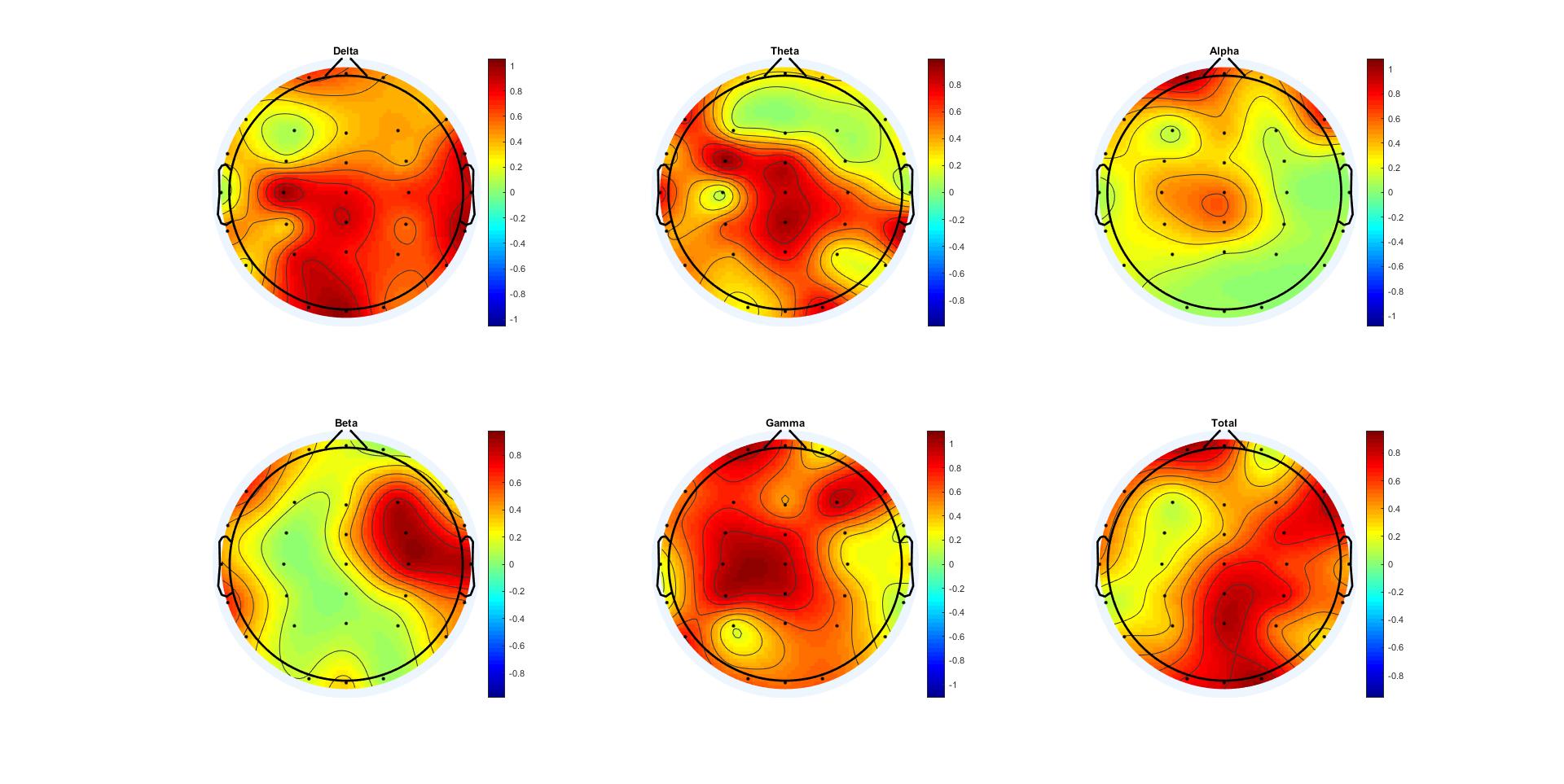

Figure8. Significant correlation between FDs of EEG signals and FDs of Peano curve animation. Colorbars indicate the Pearson's' p value.

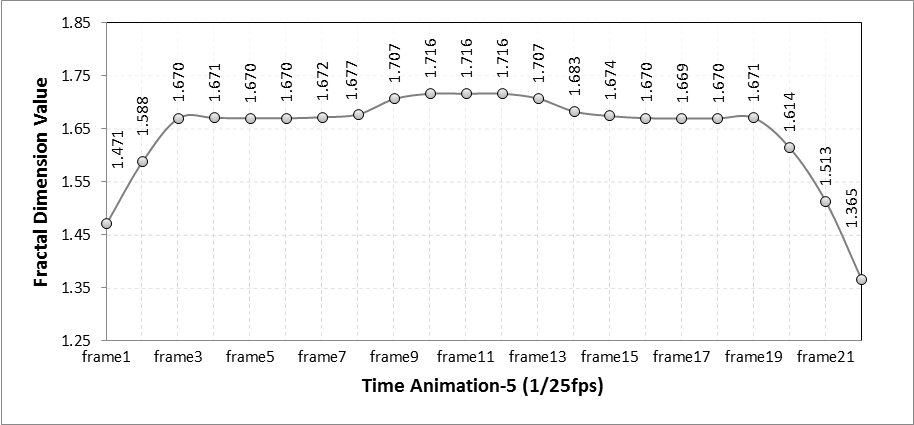

Figure9. Series of fractal dimensions of animation frames in the Pentaplexity

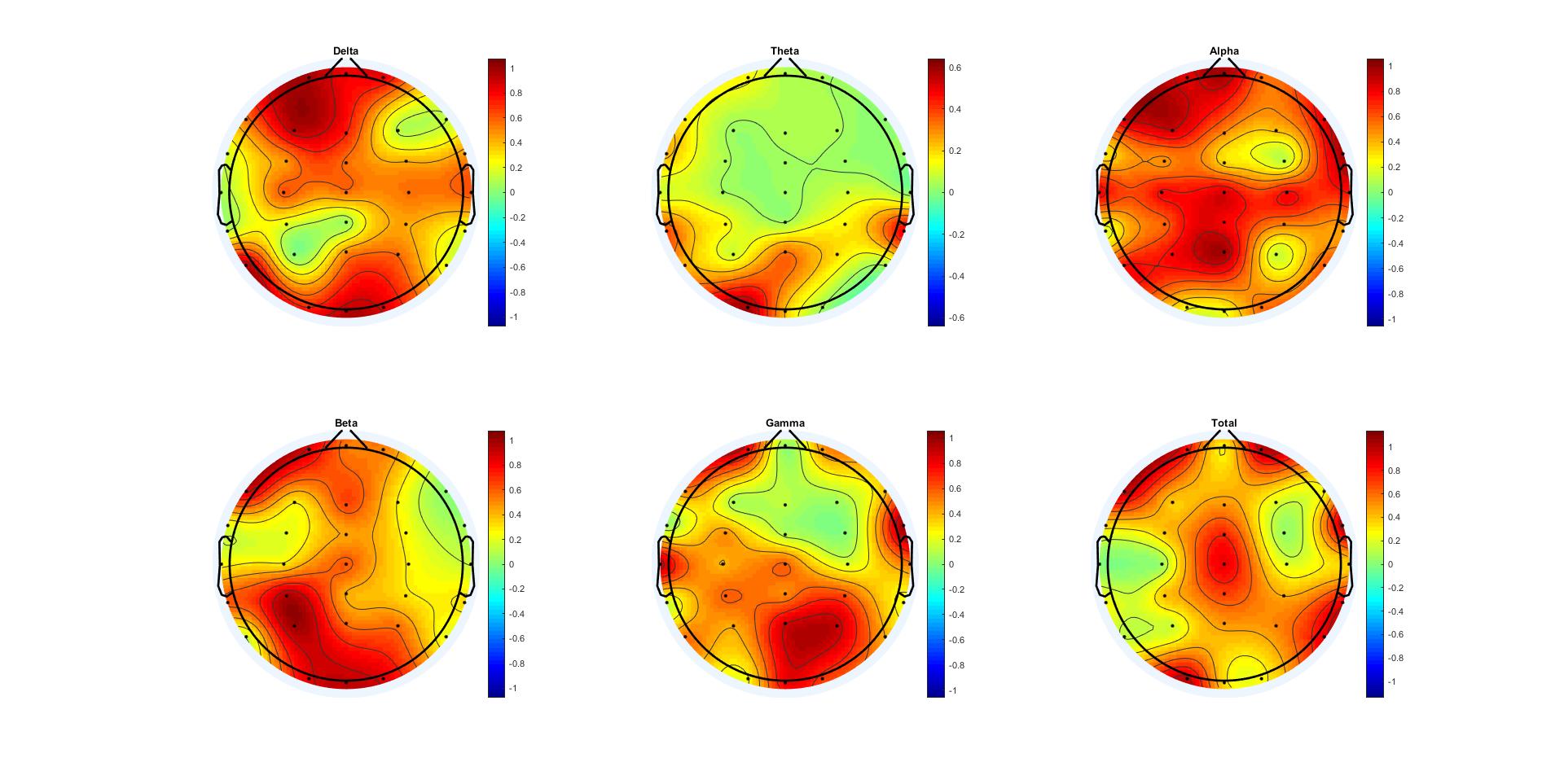

Figure10. Significant correlation between FDs of EEG signals and FDs of Pentaplexity animation. Colorbars indicate the Pearson's' p value.

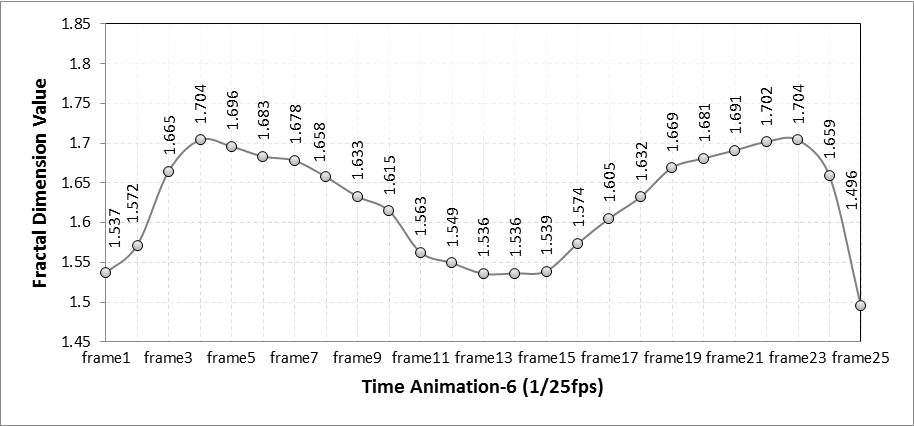

Figure11. Series of fractal dimensions of animation frames in the Penrose tiling

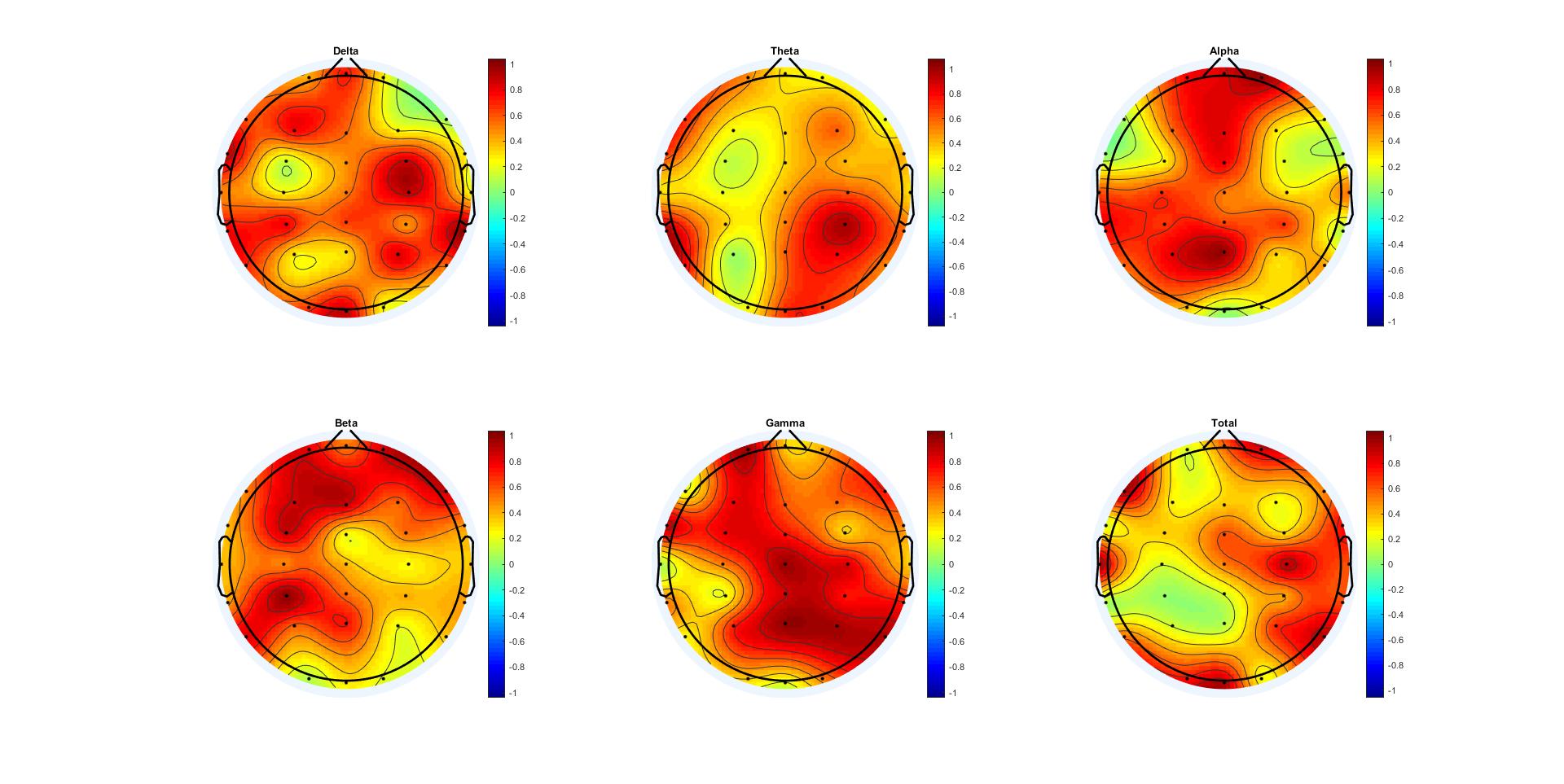

Figure12. Significant correlation between FDs of EEG signals and FDs of Penrose tiling animation. Colorbars indicate the Pearson's' p value.

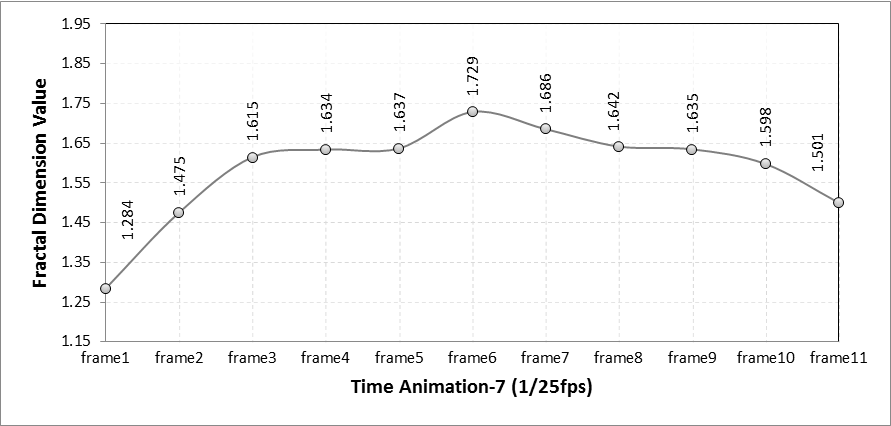

Figure13. Series of fractal dimensions of animation frames in the Dragon curve

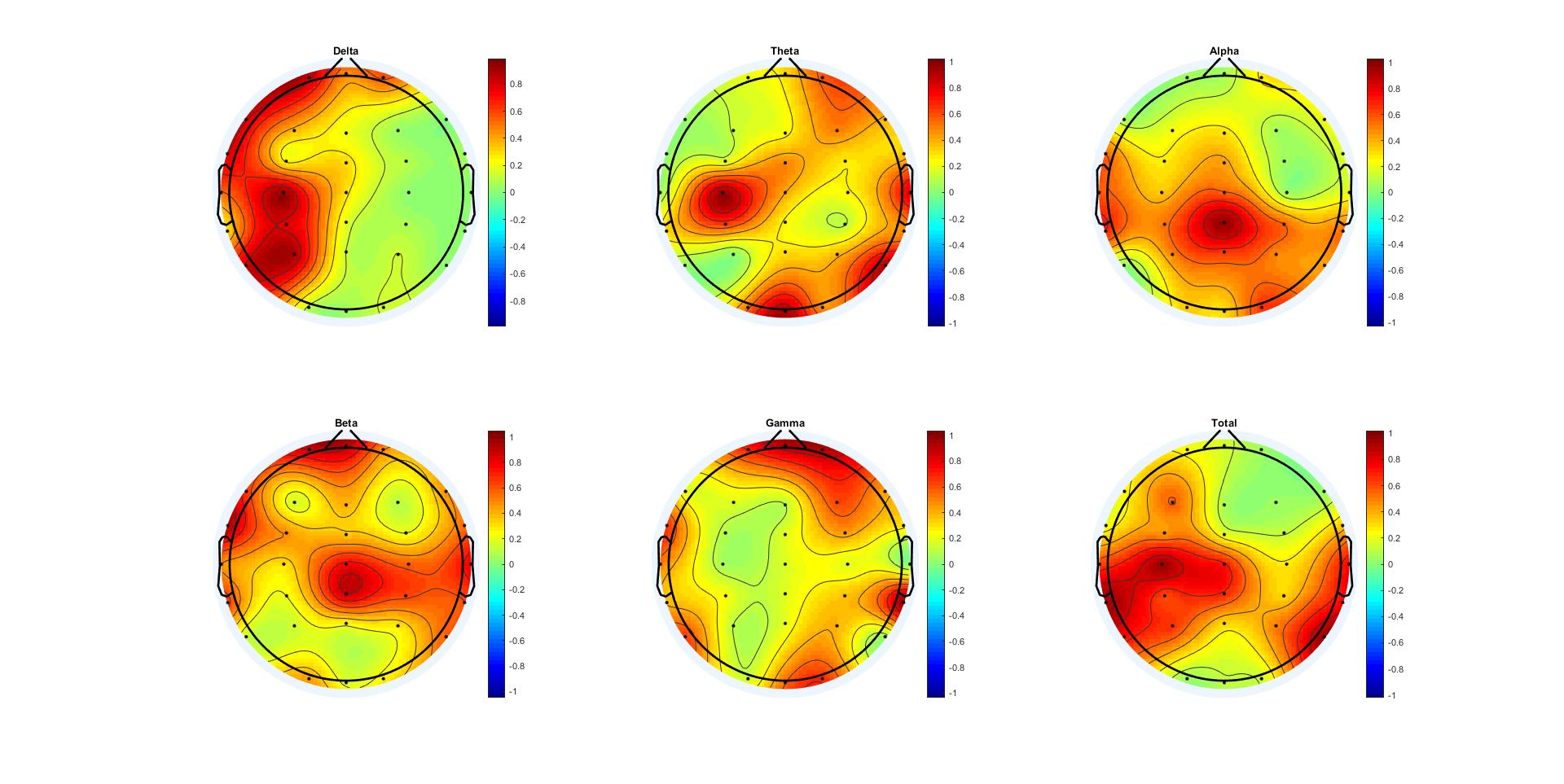

Figure14. Significant correlation between FDs of EEG signals and FDs of Dragon curve animation. Colorbars indicate the Pearson's' p value.

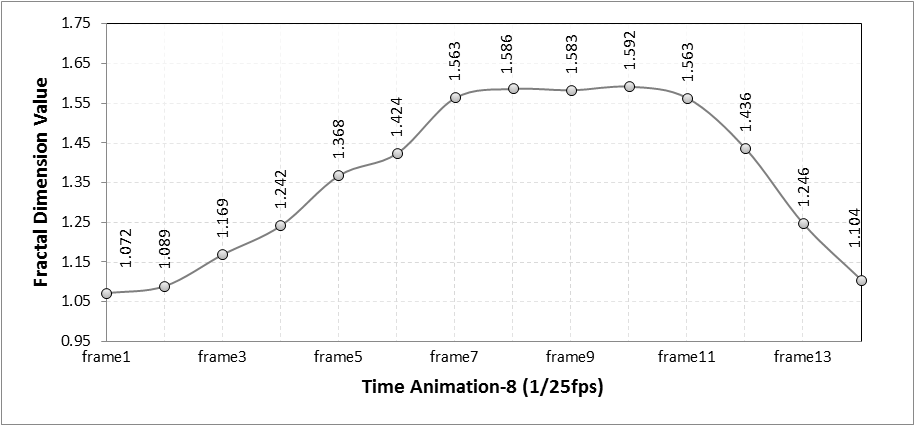

Figure15. Series of fractal dimensions of animation frames in the Terdragon curve

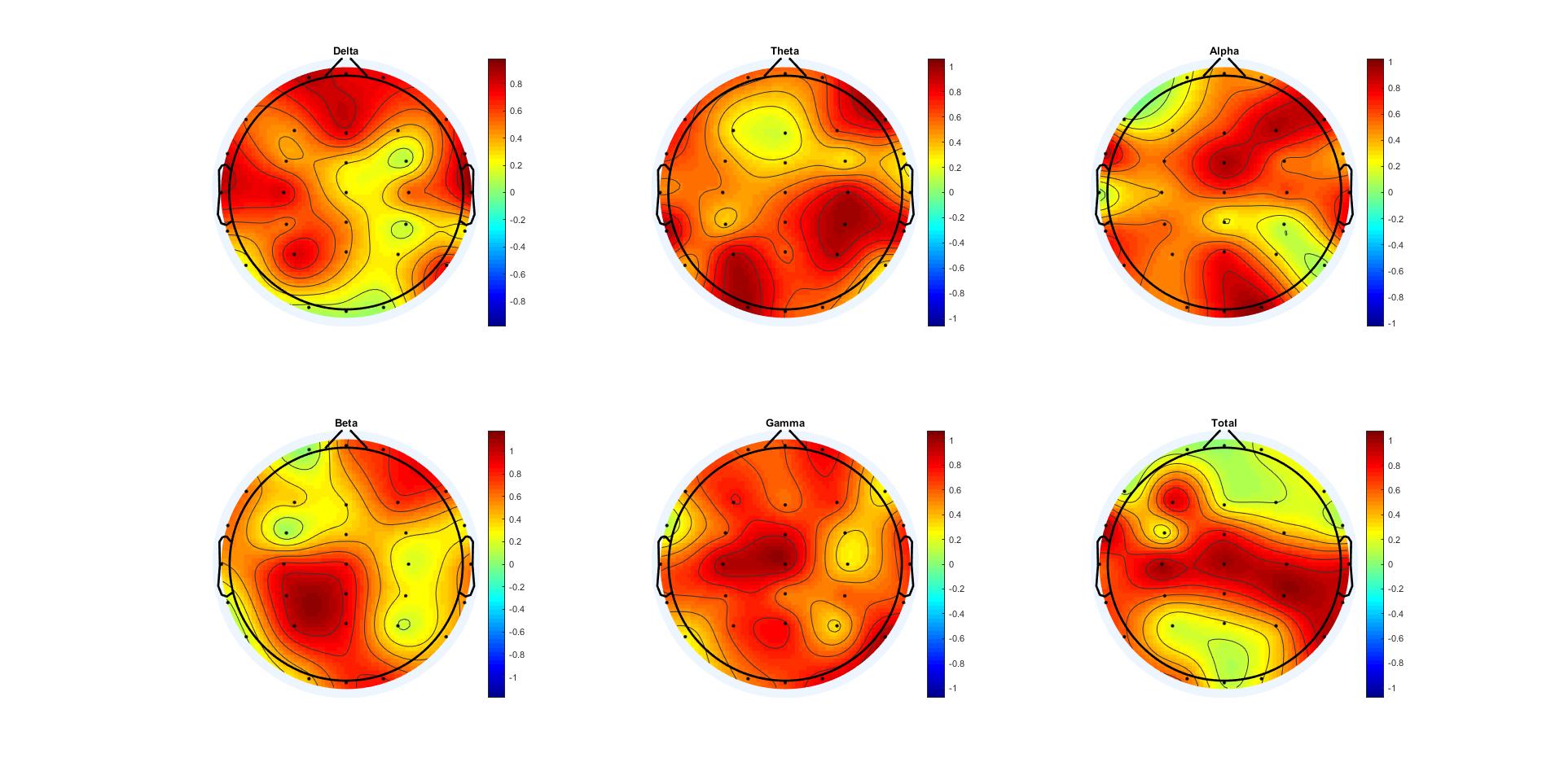

Figure16. Significant correlation between FDs of EEG signals and FDs of Terdragon curve

animation. Colorbars indicate the Pearson's' p value.

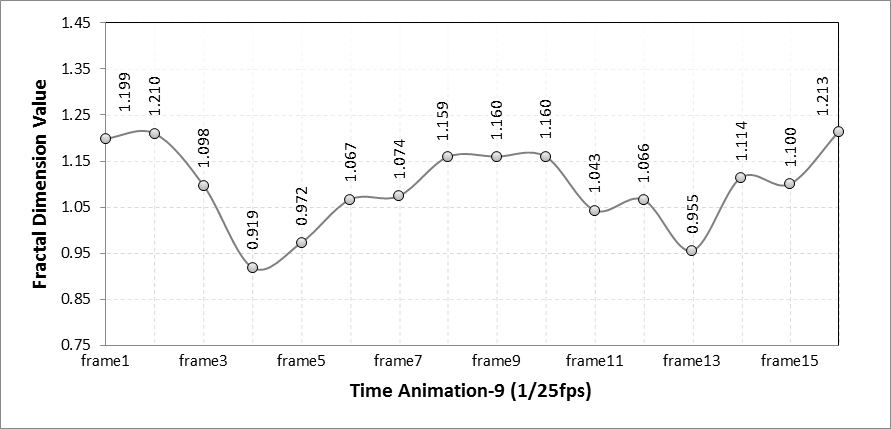

Figure17. Series of fractal dimensions of animation frames in the Lévy C curve

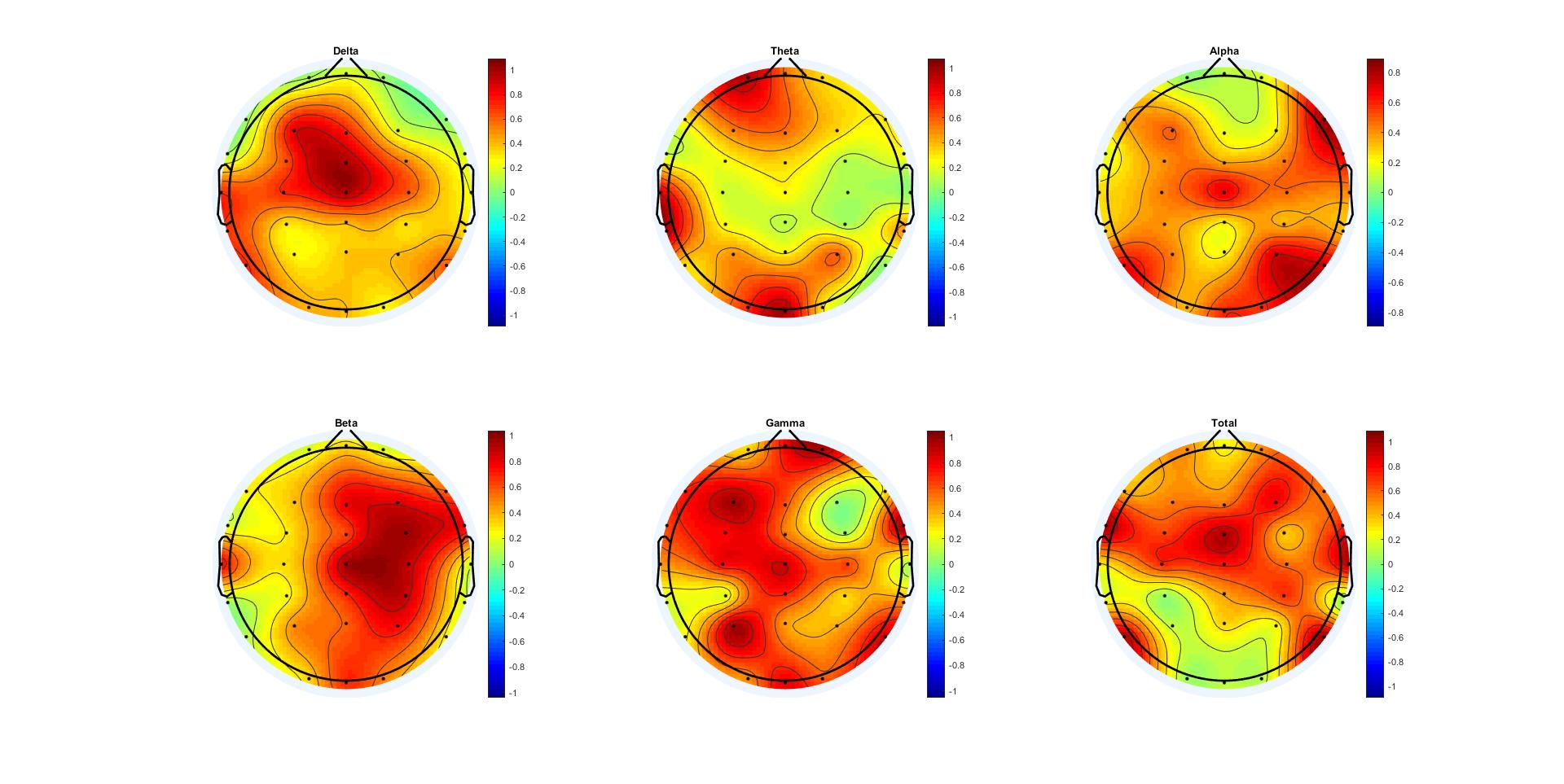

Figure18. Significant correlation between FDs of EEG signals and FDs of Lévy C curve

animation. Colorbars indicate the Pearson's' p value.

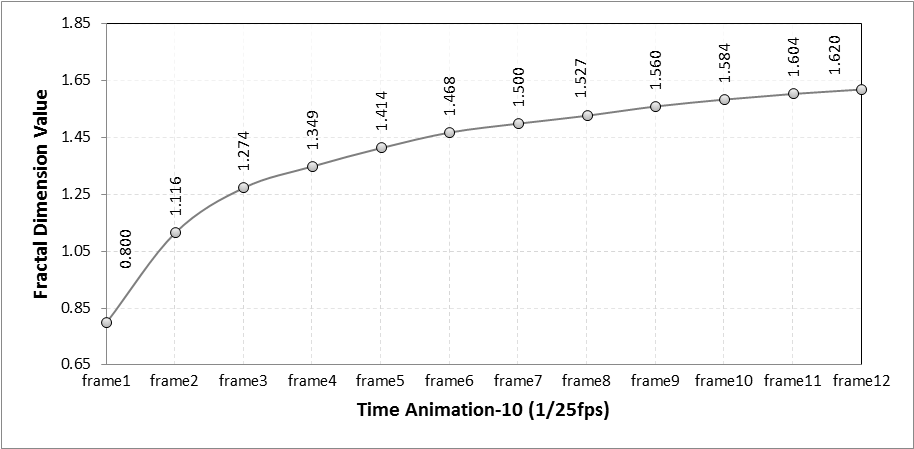

Figure19. Series of fractal dimensions of animation frames in the Multiplicative cascade

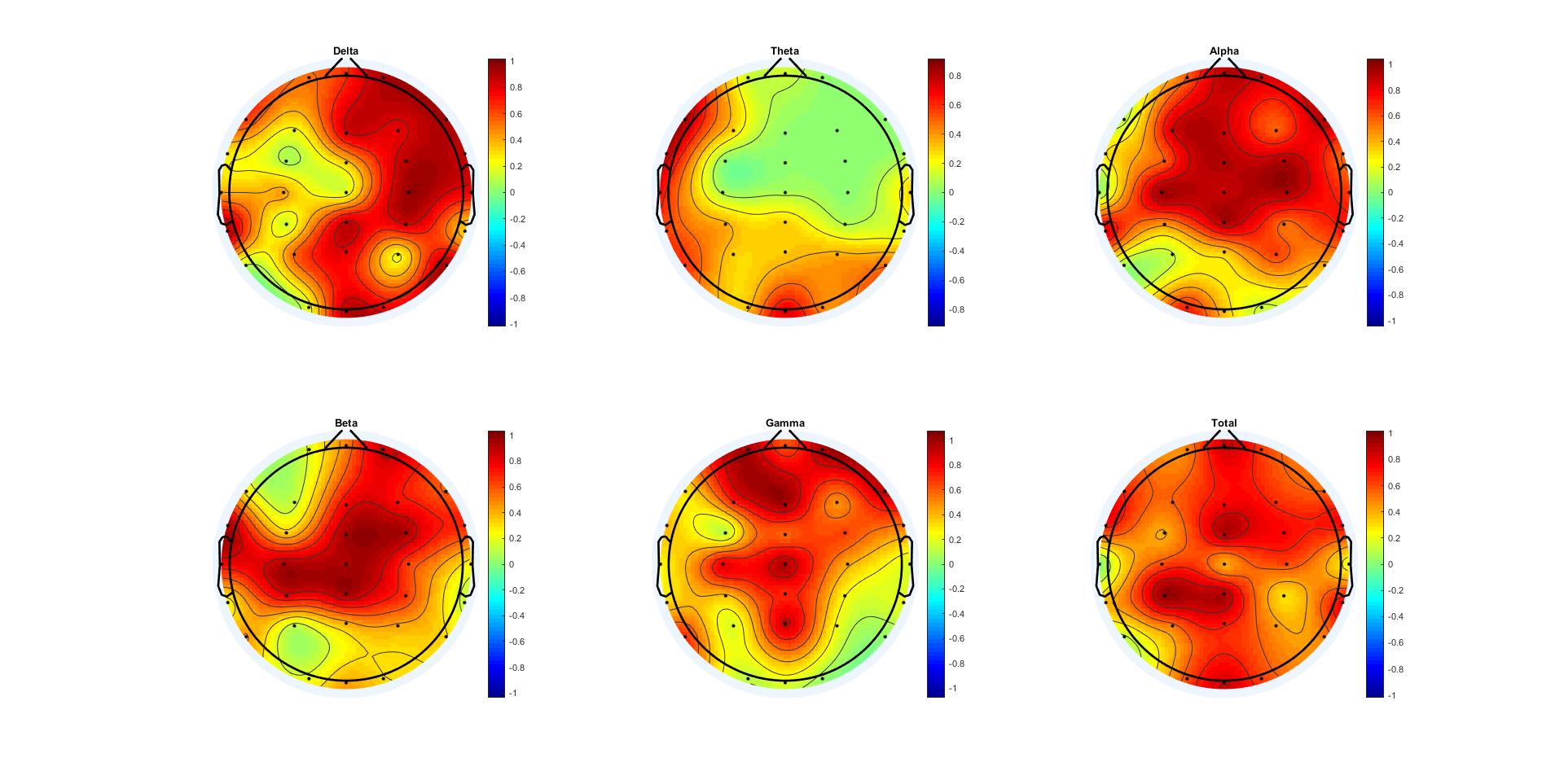

Figure20. Significant correlation between FDs of EEG signals and FDs of Multiplicative cascade animation. Colorbars indicate the Pearson's' p value.

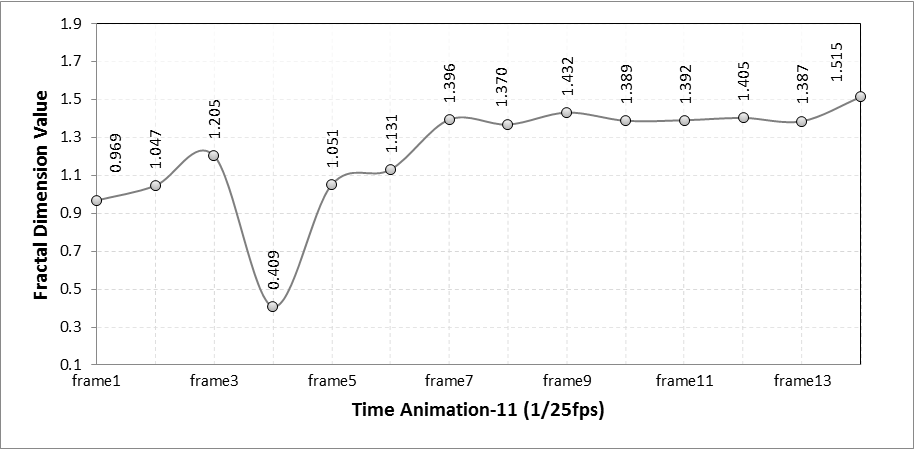

Figure21. Series of fractal dimensions of animation frames in the Diffusion-limited aggregation

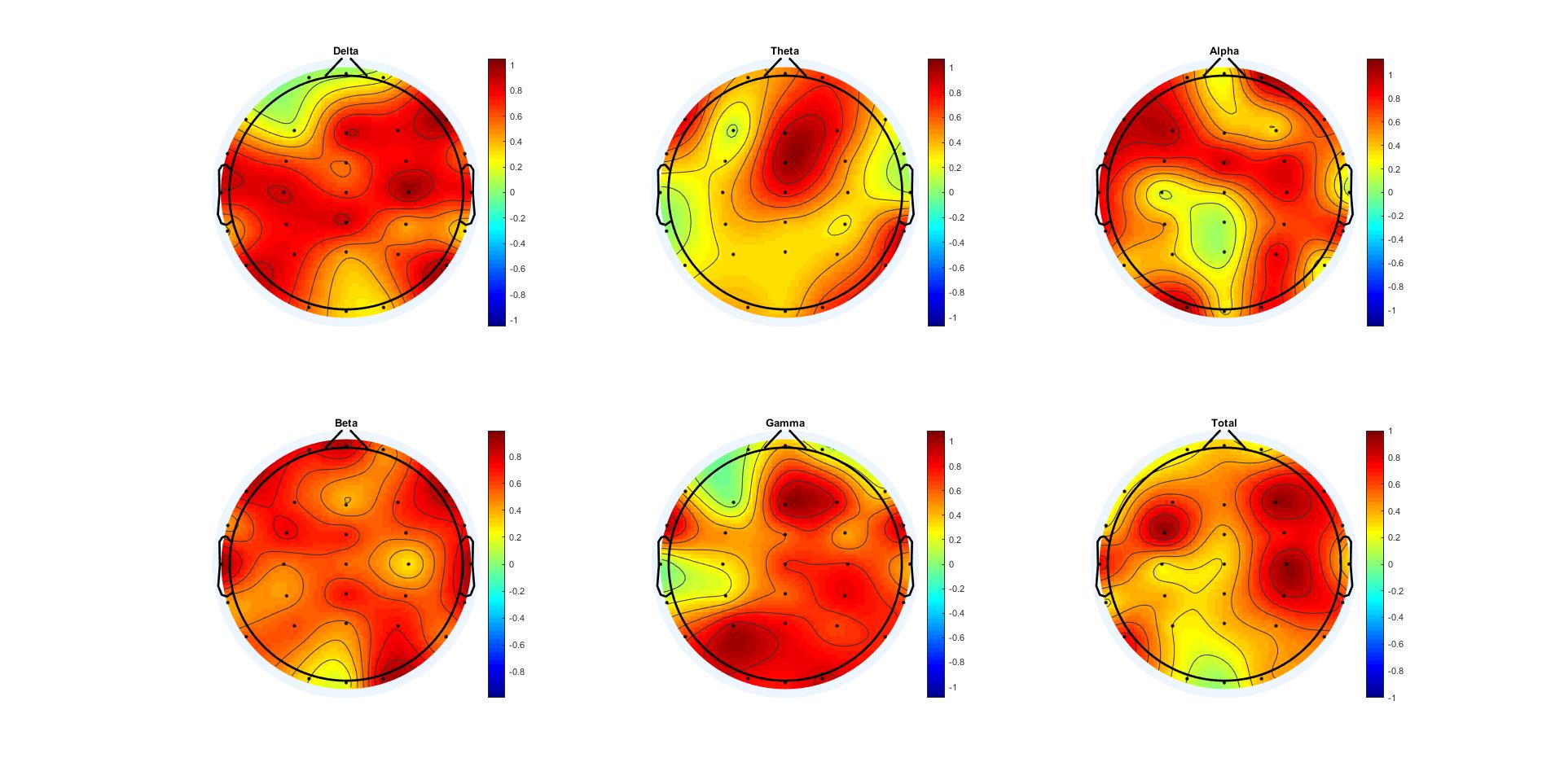

Figure22. Significant correlation between FDs of EEG signals and FDs of Diffusion-limited aggregation

animation. Colorbars indicate the Pearson's' p value.

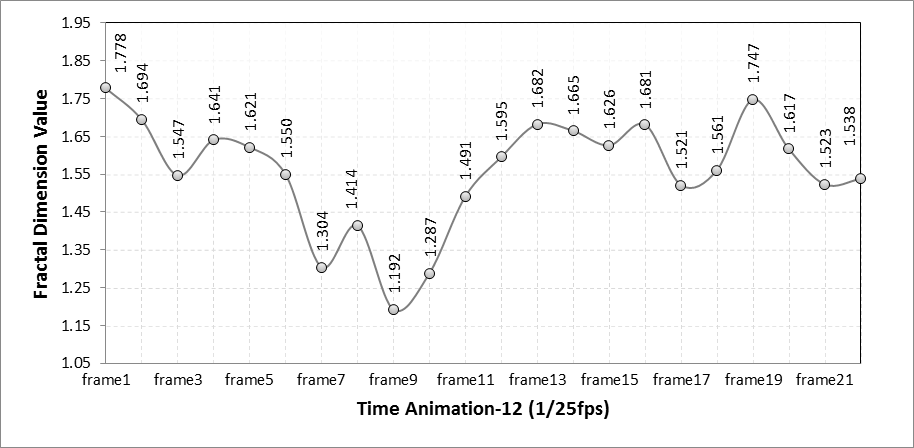

Figure23. Series of fractal dimensions of animation frames in the Mandelbrot set2

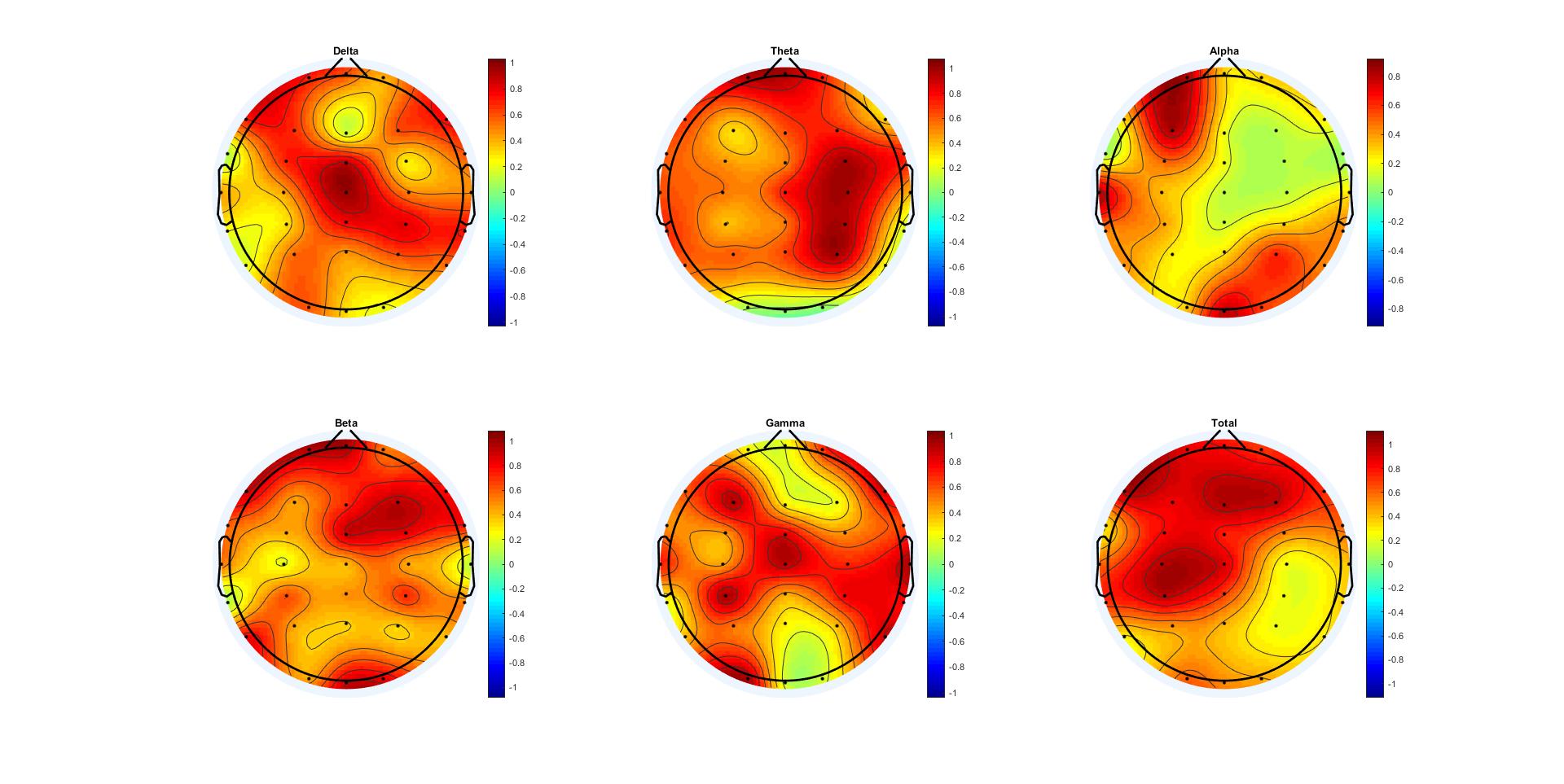

Figure24. Significant correlation between FDs of EEG signals and FDs of Mandelbrot set2 animation. Colorbars indicate the Pearson's' p value.
